# Glucose enrichment accelerates *C. elegans* reproductive aging via non-autonomous DAF-2/insulin-like receptor signaling in somatic tissues

**DOI:** 10.1101/2025.07.04.663088

**Authors:** Faria Athar, Emma J. Houston, Emily Jewett, Nicole M. Templeman

## Abstract

Detrimental effects of chronic high-sugar overconsumption can extend from molecular and cellular responses to systemic changes. Reproductive systems are particularly sensitive to diet and energetic state, yet the long-term reproductive consequences of overnutrition are poorly defined. Here, we used *Caenorhabditis elegans* to study the impacts of glucose excess on reproductive aging. Glucose supplementation shortens *C. elegans* lifespan, and we found that it also hastens age-related reproductive decline, evidenced by a greater deterioration in oocyte quality and lower fertility with age. We next evaluated insulin-like signaling contributions, as this glucose-responsive pathway is well known to regulate both somatic aging and reproductive aging. Intriguingly, while 20 mM glucose enrichment still shortens the lifespan of *daf-2(e1370)* mutants, we found that it had no detrimental impact on their reproductive aging phenotypes. Using auxin-induced tissue-selective degradation, we discovered that DAF-2/insulin-like receptor signaling in *C. elegans* intestine and body wall musculature is required for glucose enrichment to exert damaging impacts on the reproductive system. However, suppressing insulin-like signaling in either of these tissues is sufficient to protect *C. elegans* against glucose-induced reproductive aging. These findings suggest that insulin-like signalling in metabolically active somatic tissues may represent a key link between overnutrition and reproductive aging.

## Introduction

Modern diets high in energy-dense foods have led to widespread overnutrition, with physiological implications that extend beyond metabolic health. Since reproductive systems are very responsive to nutrient and metabolic cues, overnutrition has significant ramifications for reproductive health^1^—and potentially, for the age-related decline in female reproductive function. High-sugar, calorie-dense foods are associated with reduced ovarian function and subfertility, as well as greater intensity of menopausal symptoms^1–4^. It seems probable that chronic overnutrition would have cumulative impacts on reproductive aging, and metabolic conditions such as type 2 diabetes can be linked to both subfertility and earlier menopause in humans^5–9^. However, it is challenging to untangle specific mechanisms underlying these responses, because nutrient excess and metabolic dysfunction induce such breadth in changes across multiple organ systems.

The female reproductive system is one of the earliest to show age-related physiological deterioration in humans, with reductions in oocyte quality and fertility that can manifest around 30 years of age^10,11^. *Caenorhabditis elegans* is a powerful model for studying reproductive aging, with a well-defined reproductive system that also undergoes a relatively early decline driven by deteriorating oocyte quality^12^. Moreover, the *C. elegans* reproductive system is likewise attuned to metabolic cues. Evolutionarily conserved signaling systems such as insulin-like signaling respond dynamically to nutrient levels and promote energy-intensive processes like reproduction and growth, favored over preservation-type processes like stress resistance^13,14^. Suppressing insulin-like signaling by reducing levels or function of DAF-2, the only *C. elegans* insulin-like receptor, extends somatic lifespan, preserves reproductive tissue integrity with age, and significantly prolongs the fertile window. Conversely, supplementation with excess glucose generates metabolically stressful conditions for *C. elegans* that lead to shortened lifespan^15–23^ and perturbed fertility^22,24–26^. Interestingly, reduction-of-function *daf-2(e1370)* mutants are still susceptible to lifespan-shortening effects of glucose exposure^16,20^, which suggests that an assortment of metabolic responses and signaling pathways are involved in mediating glucose-induced somatic aging.

In this study, we used *C. elegans* to investigate the long-term repercussions of chronic overnutrition for reproductive aging. We found that glucose enrichment worsens oocyte quality maintenance and reduces the capacity of reproductively aged worms to produce viable progeny. Intriguingly, however, we discovered that suppressing DAF-2/insulin-like receptor signaling allows reproductive function to be preserved under glucose-enriched conditions—even if this suppression is restricted to the somatic tissues of intestine or muscle. This suggests that dysregulated insulin signaling in metabolically active somatic tissues may have important implications for the reproductive system and the progression of age-related reproductive decline.

## Results and Discussion

### Excess glucose accelerates somatic and reproductive aging

A mild dose of glucose (20 mM) was used to induce metabolically unhealthy, obesogenic-like conditions for *C. elegans* without causing excess-glucose toxicity^15^. We observed that exposure to 20 mM glucose during adulthood reduced wild-type lifespan by 15% (Fig. 1A), similar to the lifespan-shortening effects reported in other studies^15–17^. In addition to augmenting somatic aging, we found that glucose enrichment (GE) also advances age-associated reproductive decline. Reproductively aged, day 5 adult hermaphrodites were less likely to produce viable progeny when mated with young males if they had been exposed to GE during adulthood (Fig 1B). The degree of glucose impact on late-mating capacity did not appear to be dose-dependent at relatively low concentrations of 20- and 40-mM glucose added to nematode growth media (NGM).

**Fig 1.**
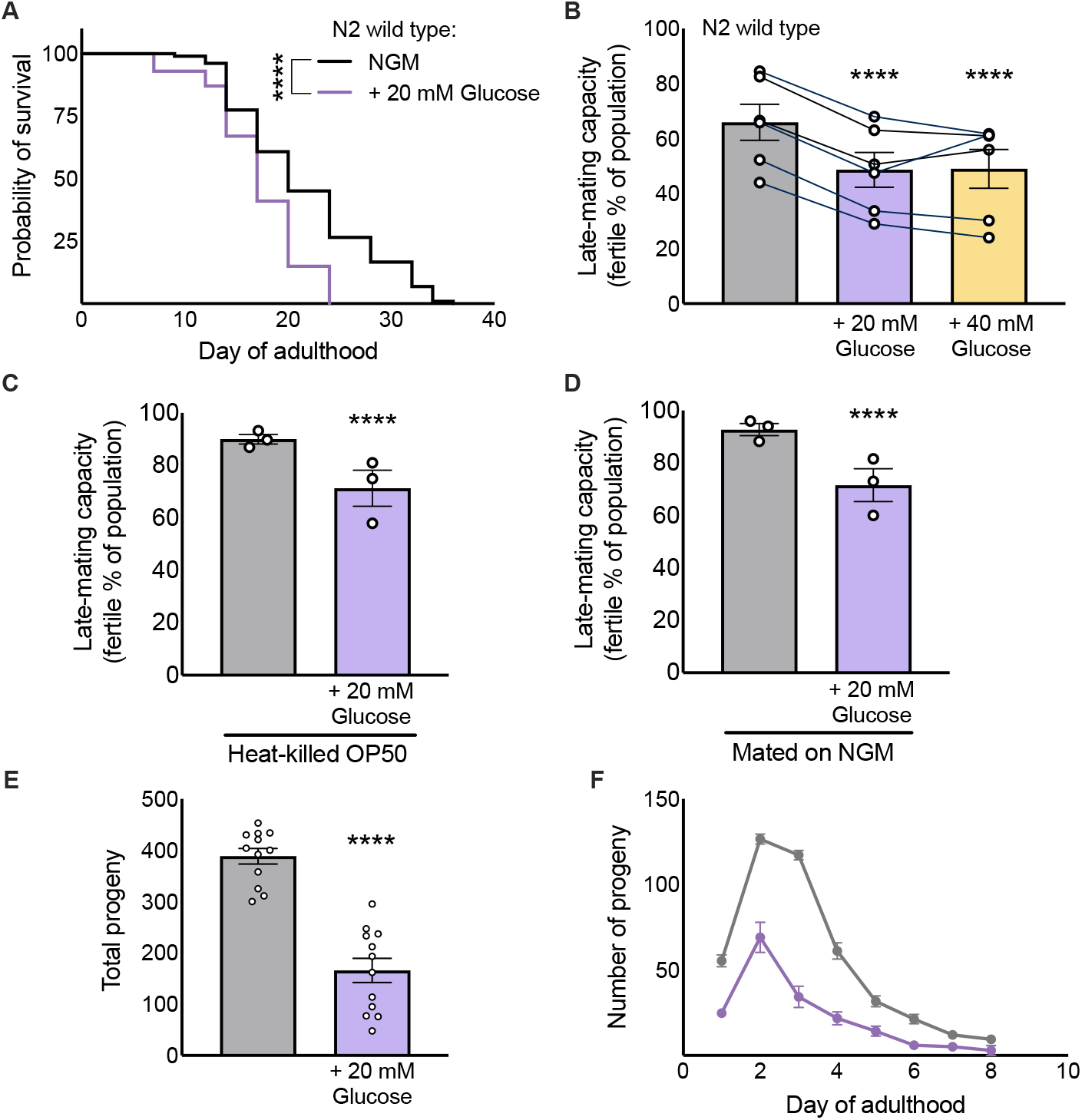
Excess glucose accelerates somatic and reproductive aging. (A) Exposure to 20 mM glucose enrichment reduces the lifespan of wildtype N2 worms compared to control NGM conditions; (n = 100 worms per group for this representative population; repeated in 2 additional independent populations; Table S1). (B) Exposure to 20 mM or 40 mM glucose decreases the late-mated reproductive capability of day 5 wild-type worms; 6 independent populations (signified by scatter points) distributed across groups, 60-130 worms per group for each population. (C, D) The effect of glucose persists when using heat-killed OP50 bacteria (C), and when all worms are mated on NGM/control plates (D). 60-100 worms per group for each population; 3 independent populations (signified by scatter plots) distributed between group. (E, F) Exposure to glucose reduces the total brood size of wildtype N2 hermaphrodites mated with *fog-2(q17)* males; scatter points signify total brood sizes for 12 individuals in a representative population; repeated in 2 additional independent populations. ∗p ≤ 0.05; ∗∗p < 0.01; ∗∗∗p < 0.001; ∗∗∗∗p < 0.0001 compared to N2 NGM. Error bars represent SEM.

Glucose has sex-specific and age-specific effects on lifespan and motility^18,19^ and can also indirectly exert impacts on worms through altering the metabolism of the OP50 *E. coli* that *C. elegans* consume^27,28^. To disentangle direct effects of glucose on worm physiology from any potential impacts of bacteria-produced glucose metabolites or other bacterial changes^27,28^, we also tested the reproductive success of day 5 adult hermaphrodites fed heat-killed bacteria throughout adulthood, and found that the detrimental effects of GE persisted without the presence of live bacteria (precluding bacteria-mediated effects; Fig. 1C). Next, we confirmed that adulthood exposure to glucose still had a significant and lasting impact when reproductively aged hermaphrodites were shifted to NGM-only plates at day 5 for the mating assay, thus ensuring that the male worms were never exposed to glucose enrichment themselves (Fig 1D). This helped us separate our observations from any potential effects of GE on males, male sperm, or mating proclivity. Glucose-exposed hermaphrodites also produced fewer progeny in total (Fig 1E,F), consistent with other published observations^22,23^. Thus, glucose exposure during adulthood directly worsens reproductive function of aging *C. elegans* hermaphrodites.

### Glucose exposure induces oocyte-level changes in worms

Oocyte quality maintenance is a key determinant of age-related reproductive decline^12,14,29^. We found that five days of glucose exposure significantly affected *C. elegans* oocyte quality, based on altered oocyte morphology and mitochondrial dynamics. Glucose-exposed worms were more likely to have maturing oocytes with an accumulation of morphological features that correspond to deteriorated quality by day 5 of adulthood, including oocytes that are abnormally small, misshapen, and/or interspersed with cavities (Fig 2A,B). Small oocyte size in worms or mammals is associated with poor quality and developmental capability of the oocyte, potentially due to maturation abnormalities or insufficient resource allocation for fertilization and embryogenesis^29–32^. The presence of cavities between neighboring oocytes or between oocytes and their surrounding somatic tissue has also been observed in older worms with poor-quality oocytes^33,34^; these morphological changes might disrupt the cellular contact required for signal transduction or molecule transfer^35^.

**Fig 2.**
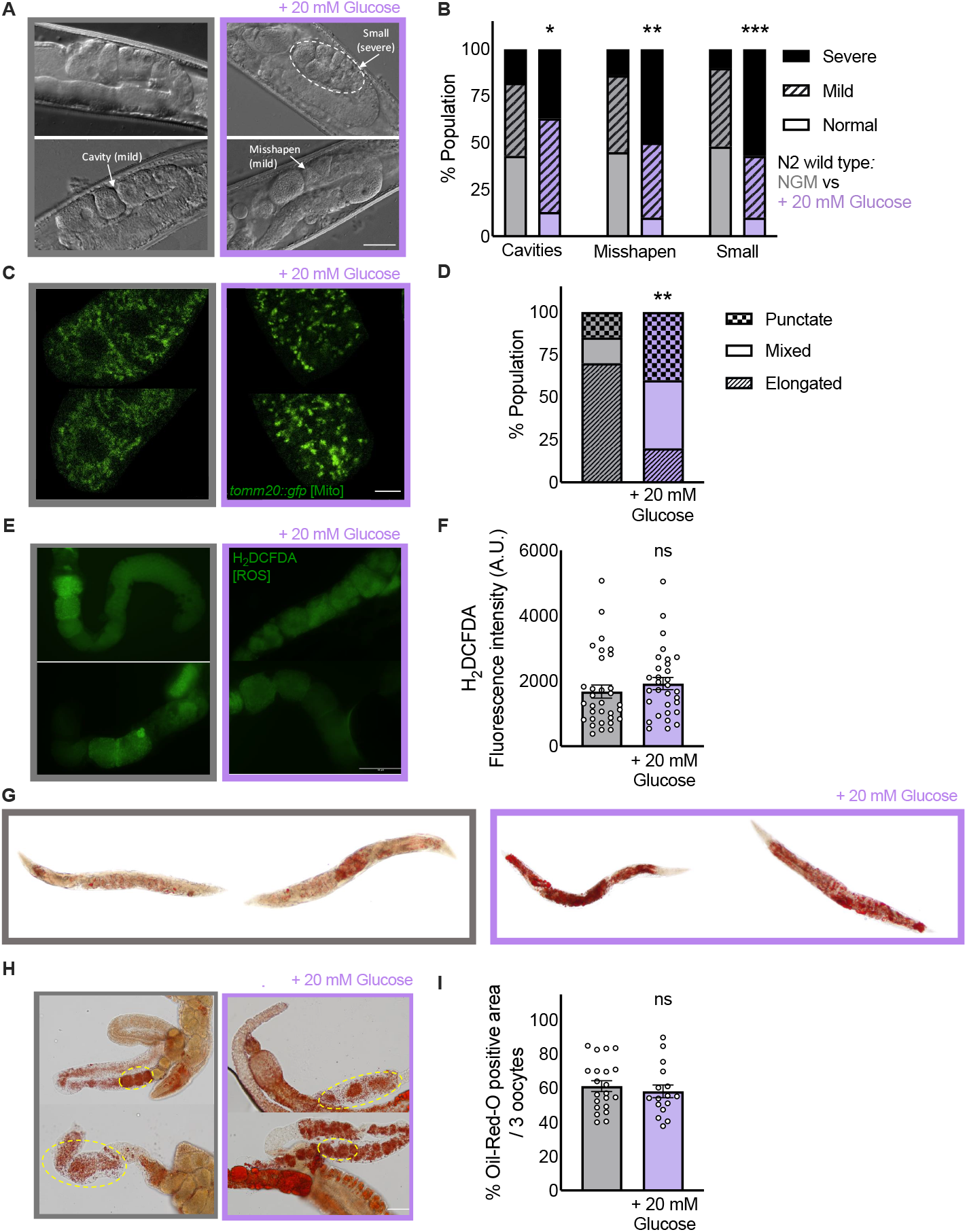
Glucose exposure alters gross oocyte morphology and oocyte mitochondrial dynamics. (A) Representative DIC images of day 5 adult wildtype oocytes, scale bar represents 25 μm. (B)Images from 29-30 worms per group were assessed for incidence of defined morphological characteristics. (C)Representative fluorescent images of the -1 oocyte of day 5 adult *mex-50:tomm-20::gfp* worms, with GFP fluorescent labelling of outer mitochondrial membranes. Scale bar represents 10 μm. (D)Images from 20 worms per group were assessed for mitochondrial morphology; mitochondria of -1 oocytes were manually scored as elongated, mixed, or punctate. (E) Representative fluorescent images of dissected gonads from day 5 adult wildtype worms dyed with H_2_DCFDA, with GFP fluorescence indicating the presence of reactive oxygen species. (F) Fluorescence intensity of the -1 oocyte Images was measured for 30 worms per group. (G) Representative images of Oil-Red-O staining of day 5 adult wild-type worms show the whole-body accumulation of lipid droplets with glucose exposure. (H) Representative color images of dissected gonads from day 2 adult wild-type worms show similar levels of lipid accumulation in oocytes irrespective of glucose exposure, scale bar represents 25 μm. (I) Oil-Red-O-positive area in the first three proximal oocytes was quantified in Images from 15-20 worms per group. ∗p ≤ 0.05; ∗∗p < 0.01; ∗∗∗p < 0.001, ns = not significant compared to N2 NGM. Error bars represent SEM, and scatter points signify data from individual worms from a representative population. All imaging experiments were performed for a minimum of 3 independent populations.

Oocytes have an abundance of mitochondria, which are involved in ATP production, calcium signaling, redox balance, and metabolism^36^. High glucose induces mitochondrial dysfunction in *C. elegans* body wall muscle^21^, thus we examined the morphology of oocyte mitochondrial networks. Cellular functions ranging from energy production to mitochondrial quality control are affected by a coordinated balance of mitochondrial fission and fusion,^37,38^ with elongated mitochondria associated with increased fusion^39^ and an increased presence of more punctate mitochondria suggesting a shift towards more fission. Using day 5 adult *mex-50:tomm-20::gfp* hermaphrodites with germline-specific labelling of mitochondria,^40^ we found that glucose-exposed worms had a higher likelihood of -1 mature oocytes with punctate or mixed mitochondrial morphology, unlike the predominantly elongated mitochondria in oocytes of age-mated NGM-exposed controls (Fig 2C,D). Imbalances in these mitochondrial dynamics can impair cellular homeostasis through such means as increasing oxidative stress, lowering mitochondrial membrane potential and respiratory capacity, and altering intracellular distribution and signaling effects^37,41^. Although glucose enrichment has been shown to promote reactive oxygen species (ROS) accumulation in whole worms^15,42^, we did not detect changed ROS levels in the oocytes of 20 mM glucose-exposed worms (Fig 2E,F). In mammals, oocytes maintain lower levels of ROS than somatic cells by limiting ROS generation in mitochondrial metabolism^43,44^; similar mechanisms might protect *C. elegans* oocytes against glucose-induced ROS accumulation.

Proper oocyte functioning is also critically dependent on normal lipid metabolism^45,46^, and GE is known to increase whole-body lipid deposition in *C. elegans*^47–50^ (see Fig 2G for examples). To gauge lipid dynamics within oocytes, we quantified lipid droplets in the three most-mature oocytes of dissected gonads, but we did not observe increased lipid content in maturing oocytes of GE-exposed, day 2 adult worms (compared to the oocytes of NGM-only controls; Fig 2H,I). Altogether, we observed that glucose enrichment compromises the maintenance of healthy oocyte morphology and mitochondrial dynamics, and these oocyte-level changes likely contribute towards the acceleration of reproductive aging.

### Reduction-of-function daf-2(e1370) mutation: protective against glucose-induced reproductive decline, but not against a shortened lifespan

Nutrient levels and energy balance is interpreted by nutrient-sensing signaling pathways that direct physiological responses. Among them, insulin-like signaling is particularly linked to glucose stimulation^16,51^. We used a *C. elegans* strain with GFP-labelled DAF-2^52^ to show that 20 mM of glucose stimulates an elevation of DAF-2 levels (Fig 3A), consistent with previous observations using a higher glucose concentration^51^. *daf-2(-)* mutants with genetic reduction of function of the insulin-like receptor are long-lived and have delayed reproductive aging,^33,34,53,54^ but *daf-2(e1370)* mutants are still susceptible to lifespan-shortening effects of glucose exposure^16,20^. Our results were consistent in showing that 20 mM GE significantly reduces the lifespan of *daf-2(e1370)* mutants (Fig 3B). Surprisingly, however, *daf-2(e1370)* worms were completely protected against a glucose-induced reduction of reproductive capacity, even exhibiting resistance to a higher, 40 mM dose of glucose. Since *daf-2(-)* mutants have longer baseline reproductive spans than wild-type worms, we performed late-mating assays on day 8 of adulthood; we found that *daf-2(e1370)* worms have similar degrees of late-mating reproductive success regardless of whether their adulthoods were spent on NGM or glucose enrichment (Fig 3C). Interestingly, when testing another reduction-of-function mutant, *daf-2(e1368)*, we found that this class I mutant was susceptible to glucose and showed similar glucose-induced reductions in late-mating capacity as wild-type worms (Fig 3D). The *daf-2(e1368)* allele contains a missense mutation in the ligand-binding domain of the DAF-2 insulin-like receptor and is associated with weaker longevity and stress resistance phenotypes than *daf-2(e1370)*, which harbors a mutation in the tyrosine kinase domain^55^. Consistent with their protection against glucose-induced reductions in late-mating success, *daf-2(e1370)* mutants also did not exhibit significant changes in brood size or oocyte quality maintenance in response to glucose exposure (Fig 3E-H). Previous studies documented reproductive decline at higher glucose concentrations (250 mM)^25^, so a higher-dose toxicity may reduce reproductive output. Collectively, these results suggest that the regulatory activities of the insulin-like receptor tyrosine kinase mediate the acceleration in reproductive aging that is caused by mild glucose enrichment.

**Fig 3.**
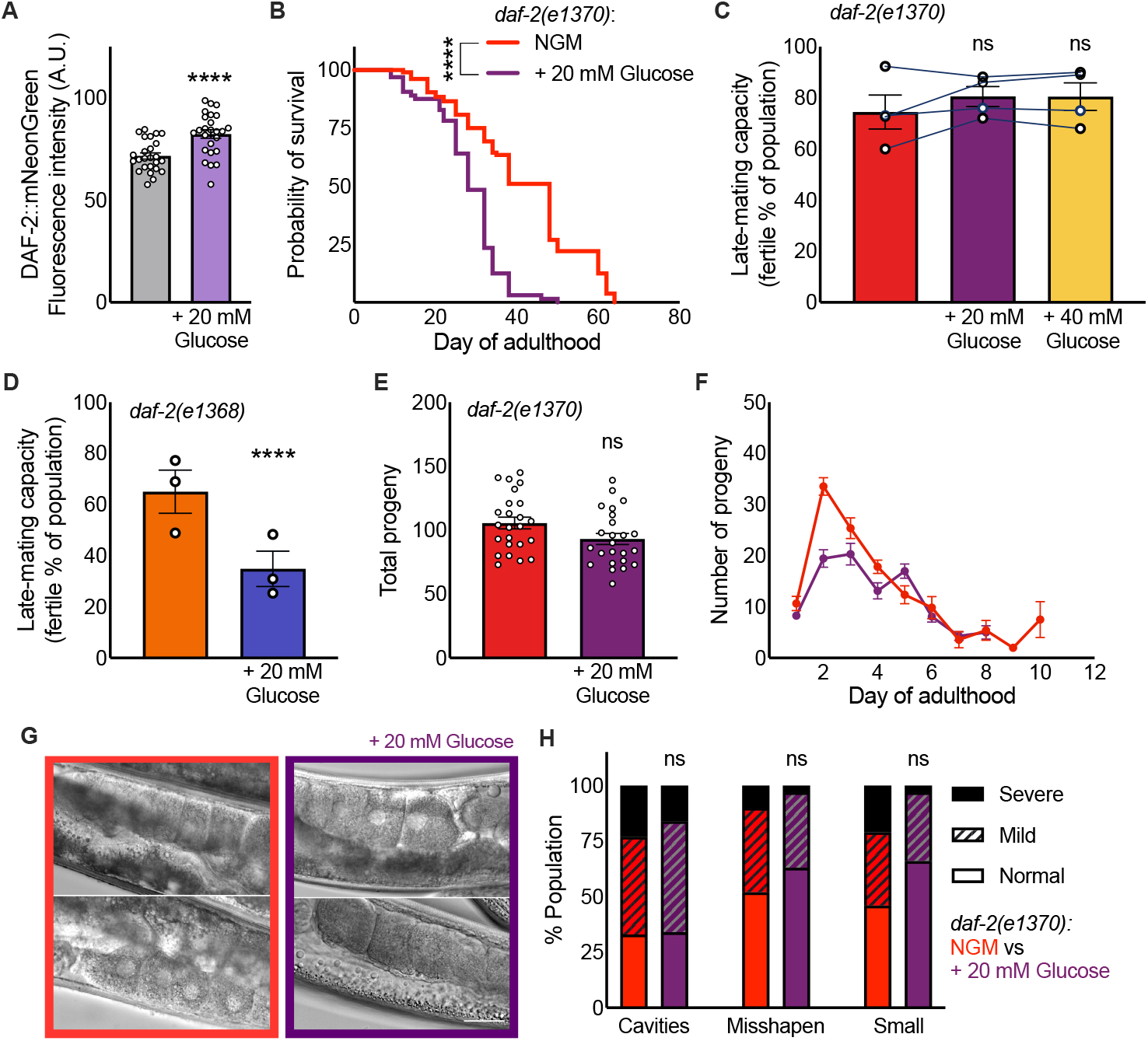
DAF-2/insulin-like receptor mediates the impact of glucose on reproductive aging but not lifespan. (A) DAF-2 levels, indicated by DAF-2::mNeonGreen, are increased on day 2 of adulthood with 20 mM glucose exposure; n = 25 worms per group (signified by scatter points) for this representative population; repeated in 2 additional independent populations. (B) Exposure to glucose during adulthood shortens *daf-2(e1370)* lifespan compared to control NGM conditions. N = 60-104 worms per group for this representative population; repeated in 2 additional independent populations (Table S1). (C) Late mated reproductive capacity of day 8 *daf-2(e1370)* remains unchanged despite 20 mM or 40 mM glucose exposure. 4 independent populations (signified by scatter points) distributed across groups, 50-104 worms per group for each population. (D) *daf-2(e1368)* shows decreased late-mating capacity with 20 mM glucose treatment; 3 independent populations (signified by scatter points) distributed across groups, 40-100 worms per group for each population. (E, F) *daf-2(e1370)* hermaphrodites have similar total brood sizes when mated with *fog-2(q17)* males despite glucose exposure, scatter plots signify total brood sizes for 24 individual worms within a single representative population; repeated in 2 additional independent populations. (G, H) Representative images and scored oocyte morphology of mated *daf-2(e1370)* worms after 8 days of glucose exposure. Images from 30-39 worms per group were analysed for this representative population; repeated in 2 additional independent populations. Error bars represent SEM; scale bar represents 25 μm.

Thus, while insulin-like signaling is important for both somatic aging and reproductive aging, it seems that this pathway might play a greater role in regulating glucose-induced reproductive system effects. This dichotomy aligns with previous observations that somatic and reproductive aging can be uncoupled through genetic manipulations. For instance, temporally restricted *daf-2* downregulation after early adulthood extends somatic lifespan without prolonging reproductive span^33,56^. Lowering *daf-*2 leads to distinct transcriptional changes in aged oocytes compared to somatic tissues, which further reinforces the concept that insulin-like signaling has diverse regulatory effects for reproductive and somatic tissues^33,34^. We show here that this compartmentalization persists under metabolically stressful conditions such as glucose enrichment, which disproportionately promotes *daf-2(e1370)* somatic aging.

### DAF-2 signaling in the intestine and body wall promotes glucose-induced reproductive decline

To further investigate the interaction between insulin-like signaling and reproductive function under glucose exposure, we used the auxin-induced degradation (AID) system for precise spatial control of DAF-2 protein levels^52^. Selective DAF-2 degradation in body wall muscle, intestine, germline, or hypodermis could improve the reproductive success of day 8 adult worms under control NGM conditions, but we found that glucose enrichment reversed that success for most strains (Fig 4A). However, glucose enrichment did not significantly worsen late-mating capacity if DAF-2 was degraded in either the intestine or the body wall musculature (Fig 4A). This suggests that the presence of insulin-like signaling in the intestine or muscle is necessary for glucose enrichment to accelerate reproductive aging. Moreover, we observed that selectively degrading DAF-2 in either of these somatic tissues—but not in the germline itself—significantly improved the late-life fertility of animals exposed to glucose (Fig 4B). Importantly, targeted DAF-2 degradation in the intestine or body wall musculature also led to improved oocyte quality under glucose enrichment, as indicated by lower frequencies of small, irregularly shaped, or cavity-interspersed oocytes (Fig 4C-F). In contrast, exposure to auxin under GE did not significantly alter oocyte quality in a germline-selective DAF-2 AID strain (Fig 4G,H), validating that these effects are non-autonomous.

**Fig 4.**
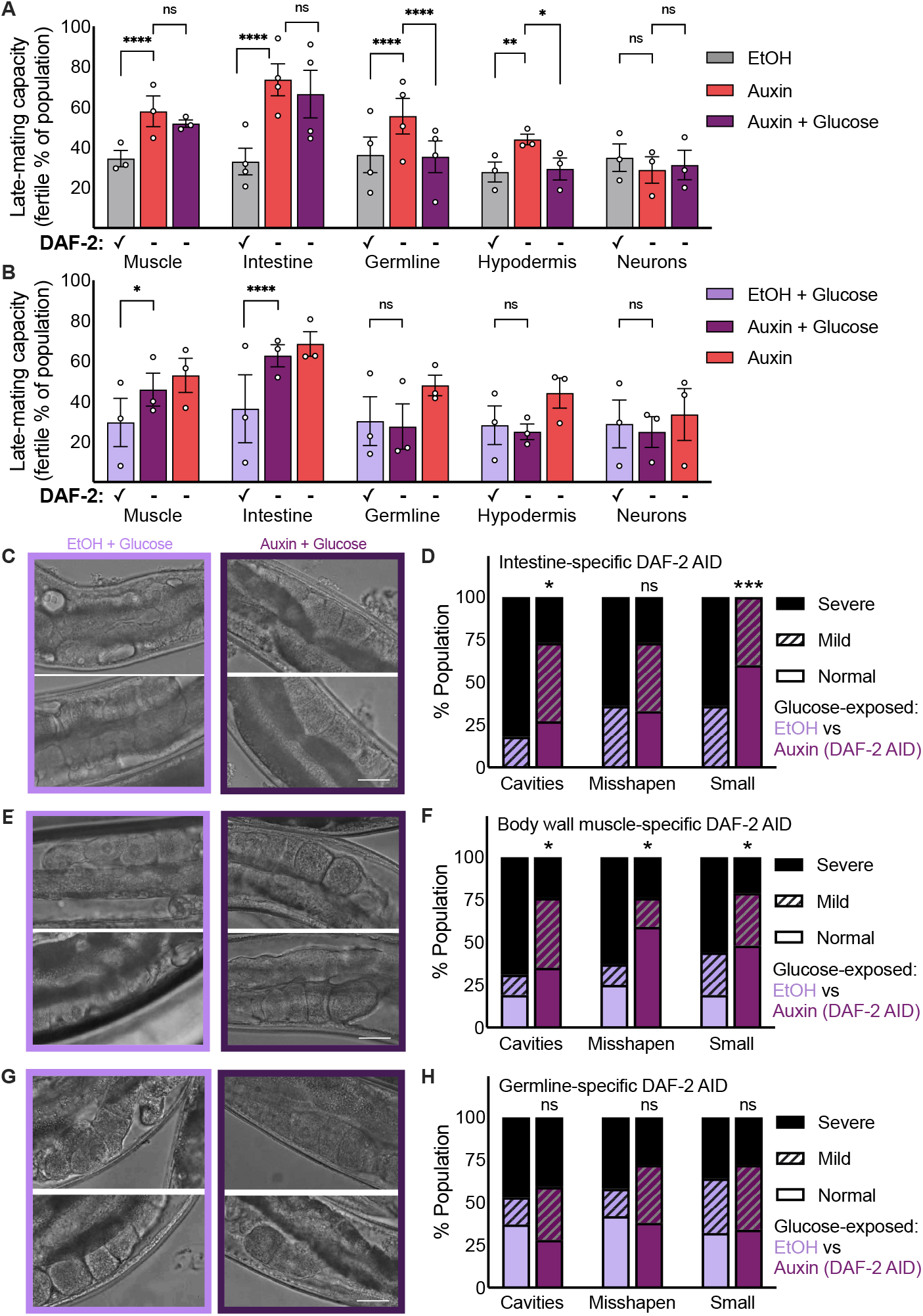
Insulin signaling in somatic tissues regulates reproductive aging under glucose stress. (A) Auxin-induced degradation (AID) of DAF-2 in body wall muscle, intestine, germline, and hypodermis (but not neurons) improves late-mating capacity of day 8 hermaphrodites under control ethanol (EtOH)-NGM conditions. Strains with AID of DAF-2 in body wall muscle, intestine, and neurons do not show glucose-induced changes in this capacity. 3 independent populations (signified by scatter points) for each strain, 30-100 worms per group for each. (B) AID of DAF-2 in body wall muscle and intestine improves late-mating capacity of glucose-exposed worms; 3 independent populations (signified by scatter points) for each strain, 30-100 worms per group for each. C, D) Intestinal DAF-2 degradation improves oocyte quality in glucose-exposed worms, as signified in (C) representative DIC images of day 8 adult oocytes, and (D) scored oocyte morphology, assessed in images from 11-15 worms per group. (E, F) Body wall muscle DAF-2 degradation similarly preserves oocyte quality in the presence of glucose, as signified in (E) representative DIC images of day 8 adult oocytes, and (F) scored oocyte morphology, assessed in images from 16-29 worms per group. (G,H). Germline DAF-2 degradation does not significantly change oocyte quality under a glucose challenge, as signified by (G) representative DIC images of day 8 adult oocytes, and (H) scored oocyte morphology, assessed in images from 16-29 worms per group. Scale bars represent 25 μm. Error bars represent SEM. ∗p ≤ 0.05; ∗∗p < 0.01; ∗∗∗p < 0.001, ns = not significant compared to N2 NGM.

Together, our data reveal that DAF-2/insulin-like receptor signaling in the intestine and body wall musculature, two metabolically and energetically important somatic tissues, promotes an acceleration of *C. elegans* reproductive aging under glucose-enriched conditions. This indicates that suppressing insulin-like signaling in these somatic tissues during glucose exposure can protect the germline against reproductive dysfunction. The insulin-like pathway DAF-16/FOXO transcription factor also acts in the intestine and muscle to extend the reproductive span of *daf-2(e1370)* mutants on NGM^33^, so these are important sites for insulin-like signaling to regulate reproductive aging across a range of metabolic conditions. The intestine has emerged as a critical regulator of reproduction in several species. For instance, in Drosophila, intestinal remodeling and metabolic rewiring directly improve fecundity and egg production^57^, and in mammals, intestinal microbiota influence hormonal balance, follicular atresia, and fertility^58^. Our findings show that the intestine can also modulate reproductive outcomes during metabolic stress in *C. elegans*, demonstrating a fundamental consequence of inter-tissue communication in response to nutritional state.

## Conclusions

In this study, *C. elegans* were exposed to glucose enrichment to explore the impacts of nutrient excess on reproductive aging. We found that glucose exposure during adulthood accelerates not only somatic aging but also reproductive aging. We also discovered that the insulin-like signaling pathway plays a crucial role in driving the glucose-induced age-related reproductive decline, through its non-autonomous actions in energetically dynamic *C. elegans* somatic tissues. Notably, reducing DAF-2/insulin-like receptor signaling prevents the glucose-induced acceleration of reproductive aging without protecting against lifespan-shortening effects, which demonstrates a regulatory dichotomy in the mechanisms by which glucose enrichment worsens reproductive system and somatic tissue functions. These findings suggest that metabolically unhealthy states associated with chronic overnutrition and upregulated somatic-tissue insulin signaling may exacerbate the age-related decline of the female reproductive system.

## Acknowledgments

We thank Alexa Fugina, Alexa Roubekas, Christian Romanowski, Ken Heschuk, Liam Wilkinson, Michel Hjelkrem, Natalie Taylor, and Taylor Sewell for their valued contributions towards performing experiments and fine-tuning experimental approaches. This study was supported by funding from the Natural Sciences Engineering Research Council of Canada (RGPIN-2022-05149). Worm strains were provided by the CGC, which is funded by NIH Office of Research Infrastructure Programs (P40 OD010440). N.M.T. is a Tier 2 Canada Research Chair in Cell Biology, a Michael Smith Health Research BC Scholar, and a member of the Sexual and Reproductive Health and Rights Research Cluster at the University of Victoria.

## Author contributions

N.M.T, F.A, and E.J.H. designed experiments and interpreted the data. F.A, E.J.H, and E.J. performed experiments and analyzed the results. F.A. and N.M.T. wrote the manuscript.

## Methods

**Table 1:**
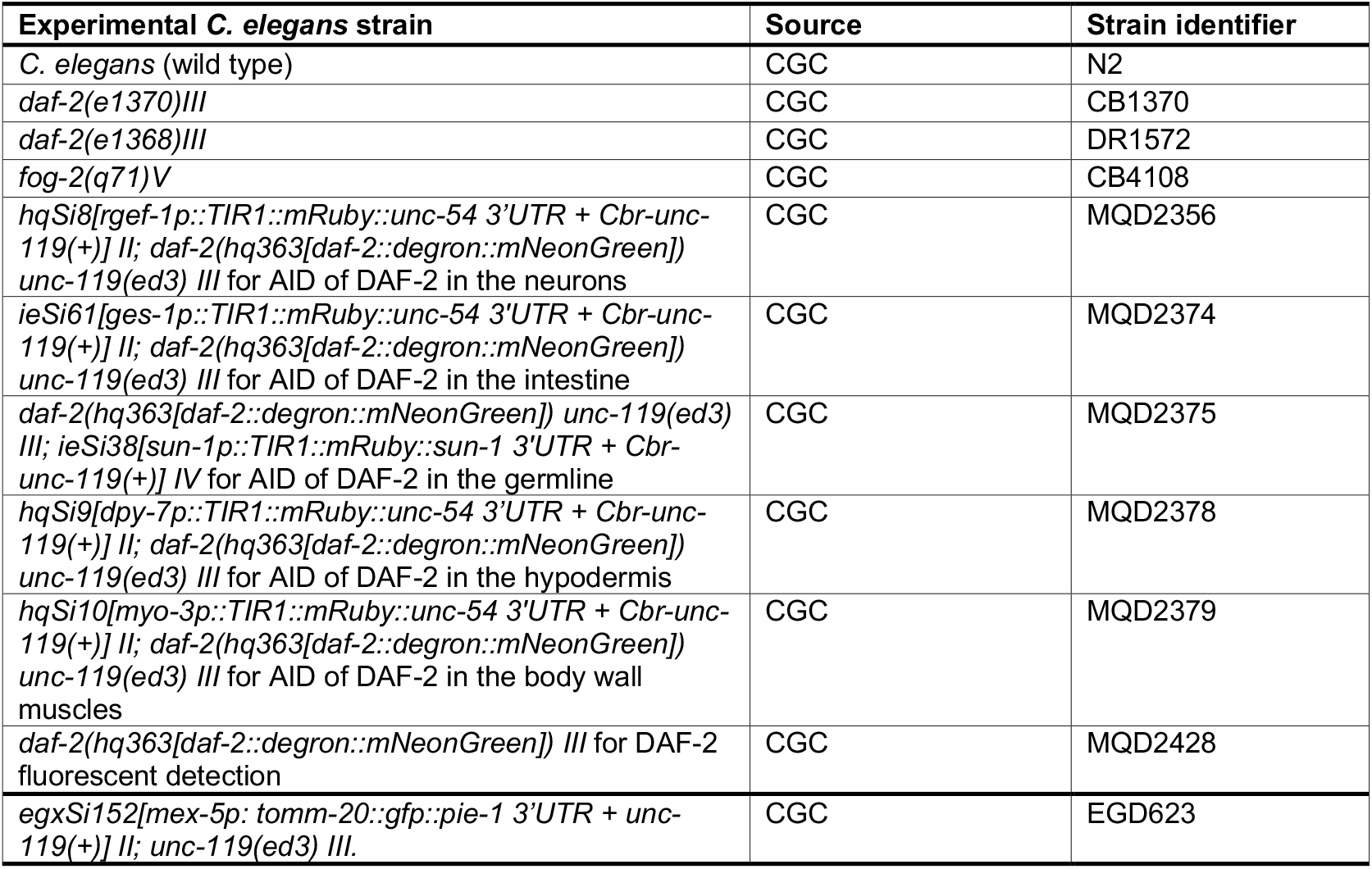
Experimental C. elegans strains.

Experiments were carried out at 20 °C on Nematode Growth Media (NGM: 3 g/L NaCl, 2.5 g/L Bacto-peptone, 17 g/L Bacto-agar in distilled water, with 1 mL/L cholesterol (5 mg/mL in ethanol), 1 mL/L 1 M CaCl2, 1 mL/L 1 M MgSO4, and 25 mL/L 1 M potassium phosphate buffer (pH 6.0)). For glucose enrichment, NGM media was supplemented with D-glucose (Sigma) to reach a final concentration of 20 mM or 40 mM in the plates. *E. coli* OP50 was used as the food source. For heat-killed OP50 experiments, overnight bacterial cultures were diluted and heat shocked at 95 °C according to a published protocol^59^. Bacterial killing was confirmed by the lack of bacterial growth on a seeded plate before use.

### Auxin-induced degradation (AID)

Auxin-containing plates were prepared to a final concentration of 1mM as previously described^52,60^. In brief, a 400 mM stock solution was prepared by dissolving auxin (indole-3-acetic acid, Sigma #I3750) in ethanol and stored at 4 °C. Auxin was added to NGM agar cooled down to 60 °C before pouring plates. Control plates for the auxin experiments had the same amount of ethanol (0.25%). Auxin plates were stored covered in aluminum foil to prevent light exposure.

### Lifespan assays

Hypochlorite-synchronized worms were transferred to glucose-enriched or NGM plates at the L4 larval stage and maintained in groups throughout their lifespans. Days of adulthood were considered to start with L4 stage as t=0 and the next day as day 1. The worms were transferred to new plates every two days during their reproductive period. If gentle prodding produced no response in an animal, it was assessed as dead. Worms were censored if they died of non-ageing related deaths such as contamination, internal hatching (bagging), or desiccation from climbing the wall of the petri plate.

### Late-mating assays

Late-mating assays were conducted as previously described^61^. Groups of hypochlorite-synchronized worms were transferred to glucose-enriched or NGM plates at the L4 larval stage and then transferred between fresh plates until they were aged to the late-mating time point, when individual hermaphrodites were distributed onto separate 35mm plates along with three day1 adult *fog-2(q71)* males per plate. Presence or absence of progeny were scored 4-5 days later; plates with dead or missing hermaphrodites or absent males were censored.

### Brood size assays

For mated brood size counts, hypochlorite-synchronized L4 larvae were individually placed on 35mm glucose-enriched or NGM plates, and young males were added in a 1:3 ratio. Mating was confirmed through the presence of male progeny, and the mated hermaphrodites were separated from males and transferred to fresh plates every 24 hours for the entirety of their reproductive span. After hermaphrodite removal, plates were kept at 20 °C and progeny was counted the next day.

### Oocyte morphology

Oocyte quality measurements were performed as previously described ^33,34^. Hypochlorite-synchronized worms were exposed to either glucose enrichment or NGM starting at the L4 stage. L4 larvae were plated at a 1:3 ratio with day 1 males for 48 hours of mating, and then successfully mated hermaphrodites (indicated by the subsequent presence of male progeny on their individual plates) were transferred between fresh plates until imaging on day 5 or day 8 of adulthood. On imaging day, worms were anesthetized with levamisole on 2% agar pads. 7-8 worms were imaged per slide on a Nikon Eclipse Ti2-E microscope using differential interference contrast (DIC) microscopy. Images were blinded and randomized prior to scoring criteria related to the phenotypes of oocyte shape, size, and connectivity. In brief, oocytes were scored as normally shaped if they were all uniformly cuboidal; mildly misshapen if there was a lack of this cuboidal shape; and severely misshapen if oocytes were visibly damaged or very irregular in shape. For the small oocyte phenotype, normal indicated that all oocyte cells filled the gonad arm and contacted the body wall; mild indicated oocytes that did not fill the gonad arm or contact the body wall; and severe indicated the presence of some miniscule oocytes. For the cavity phenotype, normal indicated no discernible cavities between oocytes; mild indicated some loss of contact between oocytes; and severe indicated the presence of distinct spaces between oocytes.

### Mitochondrial morphology assay

For the mitochondrial morphology evaluations, hypochlorite-synchronized *mex-5p:tomm-20::gfp* worms were exposed to either glucose enrichment or NGM starting at the L4 stage, when they were plated at a 1:3 ratio with day 1 *fog-2(q71)* males for 24 hours of mating. Hermaphrodites were then transferred between fresh plates every other day to prevent progeny accumulation. Upon reaching day 5 of adulthood, *mex-5p:tomm-20::gfp* worms were transferred onto 2% agar pads and fixed for 10 minutes in 3% paraformaldehyde (PFA), followed by rinsing three times with phosphate buffered saline. Oocytes were imaged using confocal microscopy with a Nikon Eclipse C2 confocal microscope. Single images and Z-stacks were taken at 60x magnification, and 2 µm slices were taken for each Z stack.

For mitochondrial analyses, images were first blinded and randomized, then the -1 oocyte of each image was scored as elongated, mixed, or punctate for mitochondrial morphology. The elongated category was indicated by oocytes with greater than half of their mitochondria being elongated, characterized by network-like structure and extensive connections with surrounding mitochondria. The mixed category was indicated by oocytes with approximately half of their mitochondria being elongated and half being punctate. The punctate category was indicated by oocytes with more than half of their mitochondria being punctate, characterized by circular morphology and a high degree of separation between mitochondria.

### ROS assay

Hypochlorite-synchronized N2 worms were placed on NGM or 20 mM glucose plates as L4 larvae and mated with day 1 *fog-2(q71)* males for 24 hours. On day 5 of adulthood, worms were anesthetized with 0.4% levamisole and gonads were dissected using two 29-gauge needles attached to 1 mL syringes. Gonads were dissected by cutting the head off the worm in a scissor-like motion just below the pharynx, which resulted in complete or partial expulsion of the gonad. Gonads were incubated in 50 nM 2’,7’-dichlorodihydrofluorescein diacetate (H_2_DCFDA) diluted in M9 for 10 minutes, followed by fixation for five minutes in 1% PFA, and rinsing three times with M9 buffer. Gonads were mounted onto 2% agar pads on microscope slides and imaged using fluorescence and DIC microscopy on a Nikon Eclipse Ti2-E at 40x magnification. Approximately 30 worms were imaged per experimental condition per independent replicate. Fluorescence intensity of the -1 oocytes (with background fluorescence subtracted) was analyzed using NIS elements software.

### Fluorescence imaging of DAF-2 levels

MQD2428 hermaphrodites with a degron and mNeonGreen tag at the 3’ end of the endogenous *daf-2* gene locus were anesthetized and placed on 2% agar pad after 2 days of adulthood spent on either 20 mM glucose-NGM or NGM alone. Fluorescence imaging was conducted on a Nikon Eclipse Ti2-E microscope at 20x magnification, and a minimum of 20-30 worms were imaged per condition. Z stacks were acquired with a slice separation of 0.6 μm, and maximal intensity projection images were analyzed using NIS elements software by subtracting mean fluorescence intensities from background fluorescence.

### Oil-Red-O staining

Mated hermaphrodites were stained with Oil-Red-O to quantify lipid presence and distribution according to the protocol by O’Rourke *et al*. ^62^. Briefly, hypochlorite-synchronized N2 worms were placed on NGM or 20 mM glucose plates as L4 larvae and mated with day 1 *fog-2(q71)* males for 48 hours. Worms were collected in M9, washed in 60% freshly prepared isopropanol, and stained with filtered Oil-Red-O solution for 18 hours. Stained worms were then washed with 0.01% Triton X-100 in S buffer and imaged using color imaging on the Nikon Eclipse Ti2-E microscope. For gonad staining, hermaphrodite gonads were dissected from day 2 adult worms as outlined in the ROS assay protocol, and were fixed in PFA before being stained with Oil-Red-O and washed with 0.01% Triton X-100 as described in^63^. ImageJ was used to quantify the spread of the red stain over the area of the first 3 proximal oocytes.

### Statistics

Lifespan assays were assessed using Kaplan-Meier survival curves (log-rank tests). Oocyte quality was assessed using Fischer’s exact tests to identify whether populations differed with respect to proportions of oocytes with severe/mild/normal phenotypes for each scored oocyte quality category. Differences in late-mating success across all replicate populations were determined using Cochran-Mantel-Haenszel tests with Bonferroni corrections applied when adjusting for multiple comparisons. Total progeny counts were evaluated with independent two-sample t-tests. Fluorescence intensities and Oil-red-O coverage were analysed using Mann-Whitney U tests. Statistical analyses were performed in GraphPad Prism and R Studio.

